# Preferred endocytosis of amyloid precursor protein from cholesterol-enriched lipid raft microdomains

**DOI:** 10.1101/2020.06.26.172874

**Authors:** Yoon Young Cho, Oh-Hoon Kwon, Sungkwon Chung

## Abstract

Amyloid precursor protein (APP) at the plasma membrane is internalized via endocytosis, and delivered to endosomes and lysosomes, where neurotoxic amyloid-β (Aβ) is produced via β-, γ-secretases. Hence, endocytosis plays a key role in the processing of APP and subsequent Aβ generation. β-, γ-secretases as well as APP are localized in cholesterol-enriched lipid raft microdomains. However, it is still unclear whether lipid rafts are the site where APP undergoes endocytosis and whether cholesterol levels affect this process. In this study, we found that localization of APP in lipid rafts was increased by elevated cholesterol level. We also showed that increasing or decreasing cholesterol levels increased or decreased APP endocytosis, respectively. When we labeled cell surface APP, APP localized in lipid rafts preferentially underwent endocytosis compared to non-raft localized APP. In addition, APP endocytosis from lipid rafts was regulated by cholesterol levels. Our results indicate that cholesterol levels regulate the localization of APP in lipid rafts affecting raft-dependent APP endocytosis. Thus, regulating the microdomain localization of APP could offer a new therapeutic strategy for Alzheimer’s disease.

## Introduction

Alzheimer’s disease (AD) is a progressive and irreversible neurodegenerative disease which is the most prevalent form of dementia (Probst et al., 1991). The hallmarks of AD pathogenesis are the extracellular deposition of senile plaques and the presence of intracellular neurofibrillary tangles, which lead to severe neuronal atrophy and ultimately death. Elevated levels of, and accumulation of cerebral β-amyloid (Aβ) peptides is the dominant pathological factor in the formation of senile plaques. Aβ is a by-product of the sequential proteolytic cleavage of amyloid precursor protein (APP) by membrane bound β-, γ-secretases (Bergmans & De Strooper, 2010, Shoji et al., 1992, Vassar et al., 1999). APP at the plasma membrane is internalized via endocytosis and delivered to early endosomes and lysosomes, where Aβ is produced. Therefore, APP is more likely to become accessible to β-, γ-secretases when the rate of APP endocytosis is increased, resulting in elevated production of Aβ (Gandy, 2005, Tanzi & Bertram, 2005). Hence, the regulation of endocytic pathways plays a key role in the trafficking and processing of APP and subsequent Aβ generation.

Cholesterol is an essential component of the plasma membrane and has number of physiological functions, among which is the regulation of endocytosis and exocytosis (Doherty & McMahon, 2009, El-Sayed & Harashima, 2013, Yue & Xu, 2015). As a result, cholesterol has been extensively implicated in the regulation of cellular APP processing, contributing to the development of AD (Burns & Rebeck, 2010, Chun et al., 2013, Grimm et al., 2017, Maulik et al., 2013, Posse de Chaves E, 2012, Urano et al., 2013, Walter & van Echten-Deckert, 2013). Furthermore, lipid dyshomeostasis, including elevated cholesterol levels, is a key participant in the pathogenesis of AD (Chun & Chung, 2020, Di Paolo & Kim, 2011, Fabelo et al., 2014, Heverin et al., 2004, Jarvik et al., 1995, Schneider et al., 2006, van Echten-Deckert & Walter, 2012, Vanmierlo et al., 2010, Xiong et al., 2008). Lipid raft microdomains, enriched with cholesterol and sphingolipids, are considered as cellular processing platforms for various cell signaling and protein-protein interactions (Brown & London, 1998, Brown & London, 2000, Pike, 2006, Sezgin et al., 2017). Substantial evidence supports the importance of lipid raft microdomains in APP processing and Aβ production. With the antibody cross-linking method, it is reported that APP and β-secretase are co-patched in lipid raft microdomains, and that Aβ production increases as cholesterol levels increase (Ehehalt et al., 2003). According to FRET experiments, clustering of APP and β-secretase in lipid raft microdomains occurs both on plasma membranes and in intracellular compartments and is triggered by exposure to cholesterol in primary neurons (Marquer et al., 2011). In addition, it is widely believed that β- and γ-secretases as well as their substrate APP are localized in lipid raft microdomains (Benjannet et al., 2001, Bhattacharyya et al., 2013, Hur et al., 2008, Osenkowski et al., 2008). Moreover, recent studies demonstrated that the enlarged lipid raft domains resulting from increased cholesterol levels enhances APP’s access to its processing enzymes, β- and γ-secretases (Cordy et al., 2003, Ehehalt et al., 2003, Marquer et al., 2011, Marquer et al., 2014, Vetrivel & Thinakaran, 2010). In accordance with these reports, a recent study reported that increased plasma membrane cholesterol induces clathrin-dependent APP endocytosis and increases Aβ generation in cultured cells (Cossec et al., 2010). Since cholesterol is the main component of lipid rafts and membrane trafficking plays a critical role in APP processing, it is important to understand the precise role of cholesterol-enriched lipid raft microdomains in APP processing and Aβ generation. In our previous study, CHO PS1 ΔE9 cells showed elevated cellular cholesterol levels that were accompanied by the increased APP localization in lipid raft microdomains (Cho et al., 2019). Modulating cellular cholesterol levels using methyl-beta-cyclodextrin (MβCD), tebuconazole, or MβCD-cholesterol redistributed APP between lipid rafts and non-rafts. These results may suggest that impaired cholesterol homeostasis is directly associated with the increased recruitment of APP into lipid raft microdomains, which then subsequently contributes to increases in Aβ production. However, it is unclear whether the increased APP localization in lipid raft microdomains is directly related to the increased Aβ production. More importantly, it is still unknown whether lipid rafts are sites where APP undergoes endocytosis.

In the present study, we found that enhanced cellular cholesterol level increase the localization of surface APP in lipid raft microdomains. Moreover, cholesterol level also regulated the rate of APP endocytosis. By labeling APP at the cell surface, we showed that APP localized in lipid rafts preferentially underwent endocytosis compared to APP in non-rafts, and that APP endocytosis from lipid rafts was regulated by cholesterol levels. The APP internalized from lipid rafts primarily accumulated in early endosomes. Furthermore, endogenous APP in hippocampal neurons was shown to relocate between lipid rafts and non-rafts depending on cholesterol levels. Thus, our findings provide compelling evidence for the involvement of cholesterol-enriched lipid raft microdomains in APP endocytosis and Aβ production.

## Results

### 1. Cholesterol levels regulate the localization of cell surface APP in lipid raft microdomains

We previously showed that cellular cholesterol level in CHO PS1 ΔE9 cells is up-regulated compared to PS1 WT cells, and that the elevated cholesterol increases APP localization in lipid rafts (Cho et al., 2019). To confirm these results, we co-immunostained surface APP with lipid raft marker, caveolin. CHO PS1 WT and PS1 ΔE9 cells were incubated with APP antibody (6E10) against the N-terminal region of APP at 4°C to label APP at the plasma membrane. After fixation, cells were permeabilized and incubated with caveolin-1 antibody. Typical immunoreactivity of APP and caveolin (cav) are shown in Fig 1A. Co-localization of APP and caveolin was quantified as shown in Fig 1B-D. The coefficient was significantly higher in PS1 ΔE9 cells compared to PS1 WT cells by 2.6-fold, indicating increased localization of APP in lipid rafts (Fig 1B, n=4). We manipulated membrane cholesterol levels with MβCD-cholesterol or MβCD to increase or decrease cellular cholesterol levels, respectively. Total cholesterol levels were measured and compared under different conditions (Appendix Figure S1). Firstly, PS1 WT cells were treated with 150 μM MβCD-cholesterol for 1 h. Cholesterol levels increased by 2.2-fold with this manipulation while APP localization in caveolin-positive lipid rafts was increased by 3-fold (Fig 1C, n=4). Secondly, PS1 ΔE9 cells were treated with 5 mM MβCD for 30 min. In this condition, we observed an approximately 27% reduction of cholesterol levels and simultaneously a significant decrease of APP localization in caveolin-positive lipid rafts (Fig 1D, n=4).

**Figure 1.**
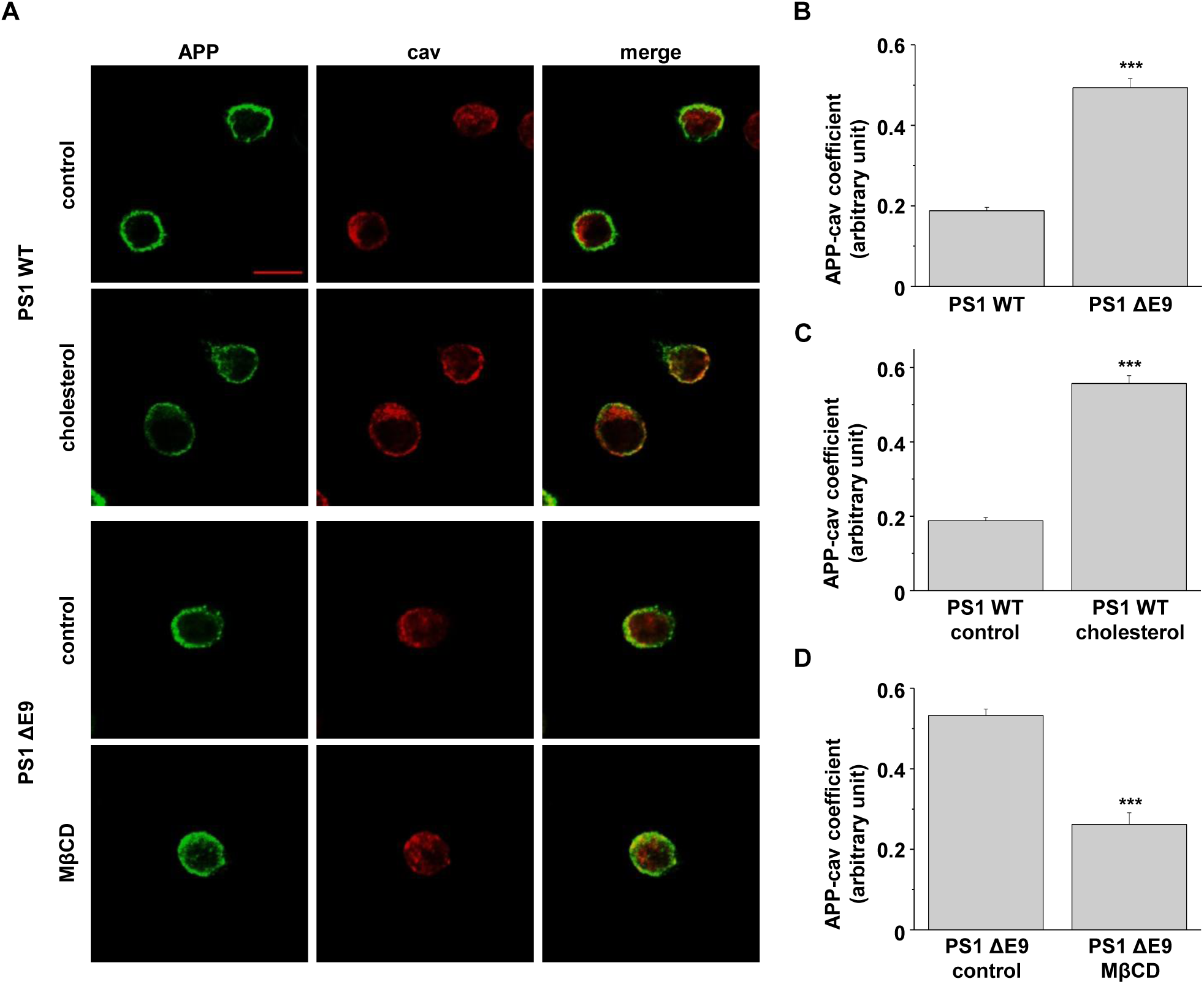
Localization of APP in lipid raft microdomains was modulated by cellular cholesterol levels. CHO PS1WT cells were incubated with 150 μM MβCD-cholesterol to increase cellular cholesterol levels. In contrast, PS1 ΔE9 cells were treated with 5 mM MβCD to reduce cellular cholesterol levels. Cells were incubated with APP antibody at 4°C to label surface APP, followed by fixing and permeabilizing. Then, cells were incubated with antibody against caveolin-1 (cav), a lipid raft marker. After washing, Alexa488- and Alexa647-conjugated secondary antibodies were used to detect primary antibodies of APP and caveolin, respectively. A. Typical immunofluorescent reactivity is displayed using confocal microscopy. Data are representative of four independent experiments. Scare bars correspond to 10 μm. B, C, D. Co-localization of APP and caveolin were indicated between (B) CHO PS1 WT cells and PS1 ΔE9 (n=4), (C) PS1 WT control and cholesterol-treated PS1 WT cells (n=4), and (D) PS1 ΔE9 and MβCD-treated PS1 ΔE9 cells (n=4). The co-localization of APP and caveolin was analyzed with Image J. Statistical analysis was analyzed by using one-way ANOVA: ***p<0.001.

Cholera toxin B (CTB) binds to ganglioside GM1, one of the components of the cholesterol-enriched lipid raft microdomains (Chinnapen et al., 2007). When we used CTB as another marker for lipid rafts, the manipulation of cholesterol levels showed similar APP localization results (Fig EV1). Taken together, these data suggested that increasing membrane cholesterol level increased APP localization in lipid rafts, while decreasing cholesterol level had the opposite effect.

### 2. Endocytosis rate of APP is increased in CHO PS1 ΔE9 cells

It is widely accepted that cellular cholesterol affects membrane trafficking, particularly through endosomal/lysosomal pathways (Doherty & McMahon, 2009, Yue & Xu, 2015). Consistent with these reports, several studies have demonstrated that cholesterol-enriched lipid raft microdomains are associated with endocytosis of membrane anchored proteins (Kirkham et al., 2005, Kirkham & Parton, 2005a, Kirkham & Parton, 2005b, Okamoto et al., 2000). Abnormal enlargement of endosomal compartments has been observed prior to the deposition of Aβ peptides in sporadic AD (Cataldo et al., 2004). Furthermore, elevated plasma membrane cholesterol increased clathrin-dependent APP endocytosis, subsequently increasing Aβ generation (Cossec et al., 2010). Since we observed that increased cholesterol levels increased the localization of APP in lipid rafts, we tested whether APP endocytosis was also affected.

For this purpose, we used the primary antibody uptake method. Two different fluorescence-conjugated secondary antibodies were used to differentiate internalized APP from APP in the plasma membrane. Cells were incubated with APP antibody at 4°C to label cell surface APP. After labeling, unbound antibodies were washed and cells were incubated at 37°C for 5, 10, and 30 min to allow internalization of antibody-bound APP. Subsequently, cells were fixed and incubated with Alexa647-conjugated secondary antibody at 4°C to visualize only the APP remaining at the plasma membrane. After permeabilizing, internalized APP was captured by Alexa488-conjugated secondary antibody. Typical immunofluorescence reactivities for surface APP (S-APP; red) and internalized APP (In-APP; green) were visualized under confocal microscopy as shown in Fig 2A. APP was exclusively localized at the plasma membrane at 0 min in both PS1 WT and PS1 ΔE9 cells. In PS1 WT cells, internalized APP gradually increased during the time course of incubation. A significant amount of APP remained at the plasma membrane even at 10 min. In PS1 ΔE9 cells, however, a significant amount of APP was internalized even at 5 min while cell surface APP was rapidly reduced. Rates of APP endocytosis were measured by calculating the intensity ratio of internalized APP (green) over surface APP (red) as shown in Fig 2B. The intensity ratio of APP at 5 min was much larger in PS1 ΔE9 cells (5.0 ± 0.3, n=5) than in PS1 WT cells (1.2 ± 0.1, n=5), indicating increased APP endocytosis.

**Figure 2.**
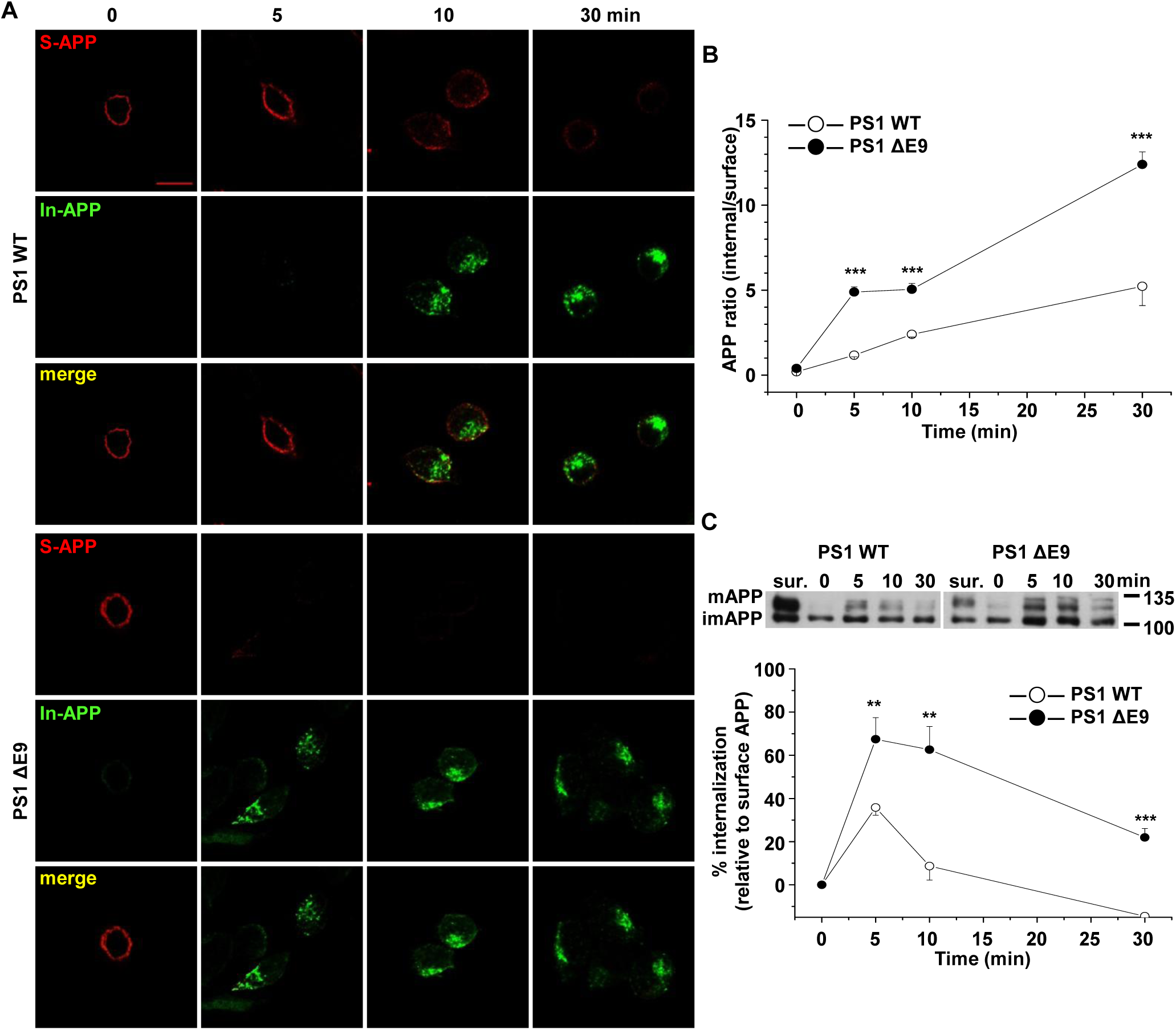
The rate of APP endocytosis was significantly increased in CHO PS1 ΔE9 cells compared to PS1 WT cells. Cells were labeled with APP antibody at 4°C and transferred to 37°C for indicated times to allow internalization. Then, cells were fixed and APP at cell surface was stained with Alexa647 (red)-conjugated secondary antibody (S-APP). After permeabilizing, Alexa488 (green)-conjugated secondary antibody was used to label internalized APP (In-APP). A. Representative confocal image shows APP localization at each time. Data are representative of five independent experiments. Scare bars correspond to 10 μm. B. Fluorescence intensities of APP were measured using Image J software. APP endocytosis was determined as the ratio of Alexa488-labeled APP/Alexa647-labeled APP (n=5). C. To biochemically quantify the rate of APP endocytosis, EZ-Link sulfo-NHS-SS-biotin was used as described in Methods. Only the internalized biotin-labeled proteins were isolated with streptavidin beads and the biotin labeled proteins were run on western blot and detected with APP antibody. Total biotin-labeled APP (surface APP; sur.) is also shown. The upper panel shows a representative western blot. The lower panel shows the rate of APP endocytosis by comparing internalized APP to surface APP (n=6). Statistical analysis was performed by one-way ANOVA: **p<0.01, ***p<0.001.

In an alternative way to quantitatively measure the rate of endocytosis, biotinylation method was used. Cells were incubated with sulfo-NHS-SS-biotin to label all surface proteins at 4°C (Ehlers, 2000, Kittler et al., 2004). After washing, cells were incubated at 37°C for 0, 5, 10, and 30 min to allow internalization of biotin-labeled surface proteins. After initiating endocytosis, reducing agent was used to remove biotins from surface proteins. Following cell lysis, the internalized biotin-bound proteins were captured using streptavidin beads. The biotin-labeled proteins were loaded on SDS-PAGE gels and examined with APP antibody. Thus, the biotin-bound APP represented the internalized APP during the incubation times at 37°C. We also labeled surface proteins with biotin and examined APP levels using Western blots to measure levels of total surface APP (surface APP). Representative Western blot image is shown in Fig 2C. The relative level of internalized APP was evaluated by comparing the amounts of biotin-labeled internalized APP to total surface APP (Fig 2C). At 5 min, 35.8 ± 3.1% of surface APP was internalized in PS1 WT cells, whereas 67.5 ± 1.0% of surface APP was internalized in PS1 ΔE9 cells (n=6). The amounts of internalized APP were significantly higher at each time point in CHO PS1 ΔE9 cells than in PS1 WT cells, consistent with our primary antibody uptake results. After 5 min, the amount of internalized APP diminished, which may be due to the rapid cleavage of APP by secretases once it is internalized (Weidemann et al., 1989). These results suggest that the endocytosis rate of APP increased in PS1 ΔE9 cells compared to PS1 WT cells.

### 3. Increased cholesterol level does not affect the endocytosis rate of transferrin in CHO PS1 ΔE9 cells

A recent study demonstrated that increasing or decreasing cellular cholesterol causes endocytosis rates to increase and decrease, respectively, in the rat calyx of Held terminals (Yue & Xu, 2015). Thus, it is possible that elevating plasma membrane cholesterol may non-specifically increase endocytosis rates of most proteins. It was also reported that cholesterol affected both clathrin-dependent and -independent endocytosis (McMahon & Boucrot, 2011, Subtil et al., 1999). To test whether the cholesterol-mediated effect was specific or non-specific, we measured transferrin endocytosis from CHO PS1 WT and PS1 ΔE9 cells. Transferrin is a well characterized protein which is internalized via clathrin-dependent endocytosis (Doherty & McMahon, 2009). Cells were treated with Alexa488-conjugated transferrin at 37°C for 5, 10, and 30 min to allow its endocytosis. The residual surface transferrin was removed by acidic buffer and the remaining intracellular transferrin was visualized under fluorescent microscopy (Appendix Figure S2). When fluorescence intensity was analyzed, the endocytosis levels of transferrin were not different between PS1 ΔE9 and PS1 WT cells. We also quantitatively measured transferrin endocytosis rate using a biochemical method. Cells were incubated with Alexa488-conjugated transferrin at 37°C for 5, 10, and 30 min to allow its internalization. Then, the cell lysates were run on Western blotting. The fluorescent bands were detected and quantified. The endocytosis rate of transferrin in PS1 ΔE9 cells was not different from that in PS1 WT cells (Appendix Figure S2). These observations suggested that the effect of increased cholesterol specifically increased APP endocytosis, but not clathrin-dependent endocytosis generally.

### 4. Cholesterol levels regulate endocytosis rates of APP

To investigate whether cholesterol levels affected the rate of APP endocytosis, we pre-treated cells with MβCD or MβCD-cholesterol. Firstly, CHO PS1 ΔE9 cells were treated with 5 mM MβCD for 30 min to reduce cellular cholesterol. Then, cells were incubated with APP antibody and further treated with two different fluorescence-conjugated secondary antibodies to visualize internalized APP and APP in the plasma membrane (Fig 3A). In MβCD-treated cells, APP remained at the plasma membrane even after 10 min internalization, which was similar to PS1 WT cells. Rates of APP endocytosis are represented by the ratio of internalized APP (green) over surface APP (red) (Fig 3B). At 5 min, the intensity ratio decreased from 7.0 ± 0.7 to 1.9 ± 0.2 (n=5) by MβCD, indicating a significant reduction of APP endocytosis. Biotinylation method using sulfo-NHS-SS-biotin was also performed to measure the rate of APP endocytosis. Cells were incubated at 37°C for 0, 5, 10, and 30 min to allow internalization (Fig 3C). Levels of total surface APP (surface APP) were also measured. In PS1 ΔE9 control cells, 73.2 ± 9.7% (n=6) of surface APP was internalized at 5 min. In contrast, only 26.8 ± 5.7% (n=6) of surface APP was internalized at 5 min in MβCD-treated cells. These results indicate that decreasing cellular cholesterol levels in PS1 ΔE9 cells decreased APP endocytosis rate.

**Figure 3.**
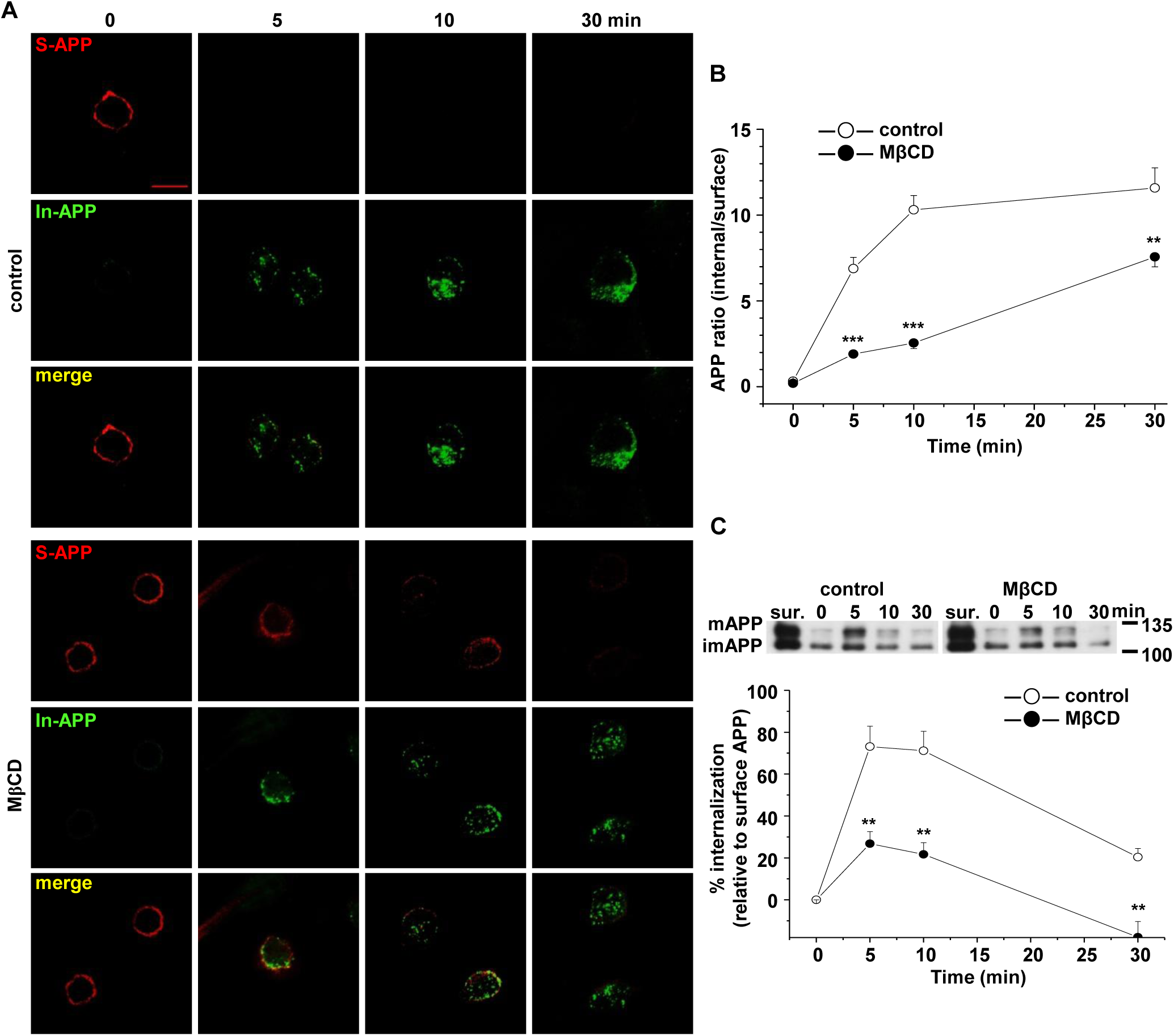
Reduction of cellular cholesterol decreased the rate of APP endocytosis in PS1 ΔE9 cells. CHO PS1 ΔE9 cells were treated with 5 mM MβCD. Cells were then labeled with APP antibody at 4°C to monitor APP endocytosis as described in Fig 2. A. Representative confocal image shows APP localization at indicated times. Data are representative of five independent experiments. Scare bars correspond to 10 μm. B. The rate of APP endocytosis was measured as the ratio of internalized APP over surface APP (n=5). C. APP endocytosis was quantified using EZ-Link sulfo-NHS-SS-biotin as described in Fig 2. Total biotinylated APP (surface APP; sur.) is also shown. The upper panel shows a representative western blot. The lower panel shows the rate of APP endocytosis by comparing internalized APP to surface APP (n=6). Statistical analysis was carried out by one-way ANOVA: **p<0.01, ***p<0.001.

Next, we exogenously elevated cellular cholesterol levels by treating CHO PS1 WT cells with 150 μM MβCD-cholesterol for 1 h. The rate of APP endocytosis was measured with primary antibody uptake method. In PS1 WT control cells, a large amount of surface APP remained at the plasma membrane at 10 min (Fig EV2). In contrast, surface APP was reduced significantly even at 5 min in MβCD-cholesterol-treated cells. The ratio of APP endocytosis was 1.9 ± 0.1 (n=4) and 3.4 ± 0.2 (n=4) in PS1 WT control cells at 5 min and 10 min, respectively. MβCD-cholesterol increased the ratio of APP endocytosis to 9.2 ± 1.4 (n=4) and 12.0 ± 1.2 (n=4), respectively. Thus, increasing cellular cholesterol levels in PS1 WT cells increased the rate of APP endocytosis. Taken together, these results suggested that cholesterol levels regulate APP endocytosis.

### 5. APP localization in lipid rafts is regulated by cholesterol levels

In our previous report, we showed that localization of APP in lipid rafts is increased in CHO PS1 ΔE9 cells compared to PS1 WT cells (Cho et al., 2019). We also showed that there is a direct correlation between cellular cholesterol levels and APP localization in lipid raft fractions. To quantitatively measure APP localization in lipid raft microdomains, we designed a biotin-labeled lipid raft fractionation method. Cells were pre-treated with EZ-Link NHS-biotin at 4°C for 10 min to label cell surface proteins. After unbound biotins were washed, cells were harvested, homogenized, and sonicated. Equal amounts of biotin-bound proteins from cell lysates were loaded on discontinuous sucrose density gradients as described in Methods. Equal volumes from 12 recovered gradient fractions were run on Western blot to detect APP, β-actin (a non-lipid raft marker), and caveolin (a lipid raft marker) as shown for a typical result in Fig EV3. When levels of cholesterol and protein were measured, cholesterol-enriched lipid raft fractions (4 to 6) were clearly separated from protein-enriched non-raft fractions (8 to 12). Protein levels in each fraction did not differ between the PS1 WT and PS1 ΔE9 cells. However, cholesterol levels in lipid raft fractions were significantly higher in PS1 ΔE9 cells than in PS1 WT cells as we reported previously (Cho et al., 2019). For better quantification of APP levels in lipid raft and non-raft fractions, we combined fractions from 4 to 6 (lipid raft fractions, R), and fractions from 8 to 12 (non-raft fractions, NR). Equal amounts of biotin-bound surface proteins were captured with streptavidin beads. Then, equal volumes of each fraction were loaded on gels for Western blots. Thus, the biotin-bound APP represented cell surface APP localized either in lipid raft or in non-raft fractions. A typical result is shown in Fig 4A and 4C. Higher APP levels in lipid raft fractions compared to non-raft fractions were due to significantly lower total protein levels in lipid raft than in non-raft fractions. Thus, APP comprised a larger proportion of the total proteins in lipid raft fractions. The ratio of surface APP levels in each fraction was calculated as shown in Fig 4B and 4D. APP level in raft fractions was higher in PS1 ΔE9 cells (93.2 ± 4.0%, n=5) than in PS1 WT cells (71.6 ± 4.6%, n=6), which was consistent with our previous co-staining result in Fig 1. Conversely, surface APP level in non-raft fractions was lower in PS1 ΔE9 cells compared to PS1 WT cells.

**Figure 4.**
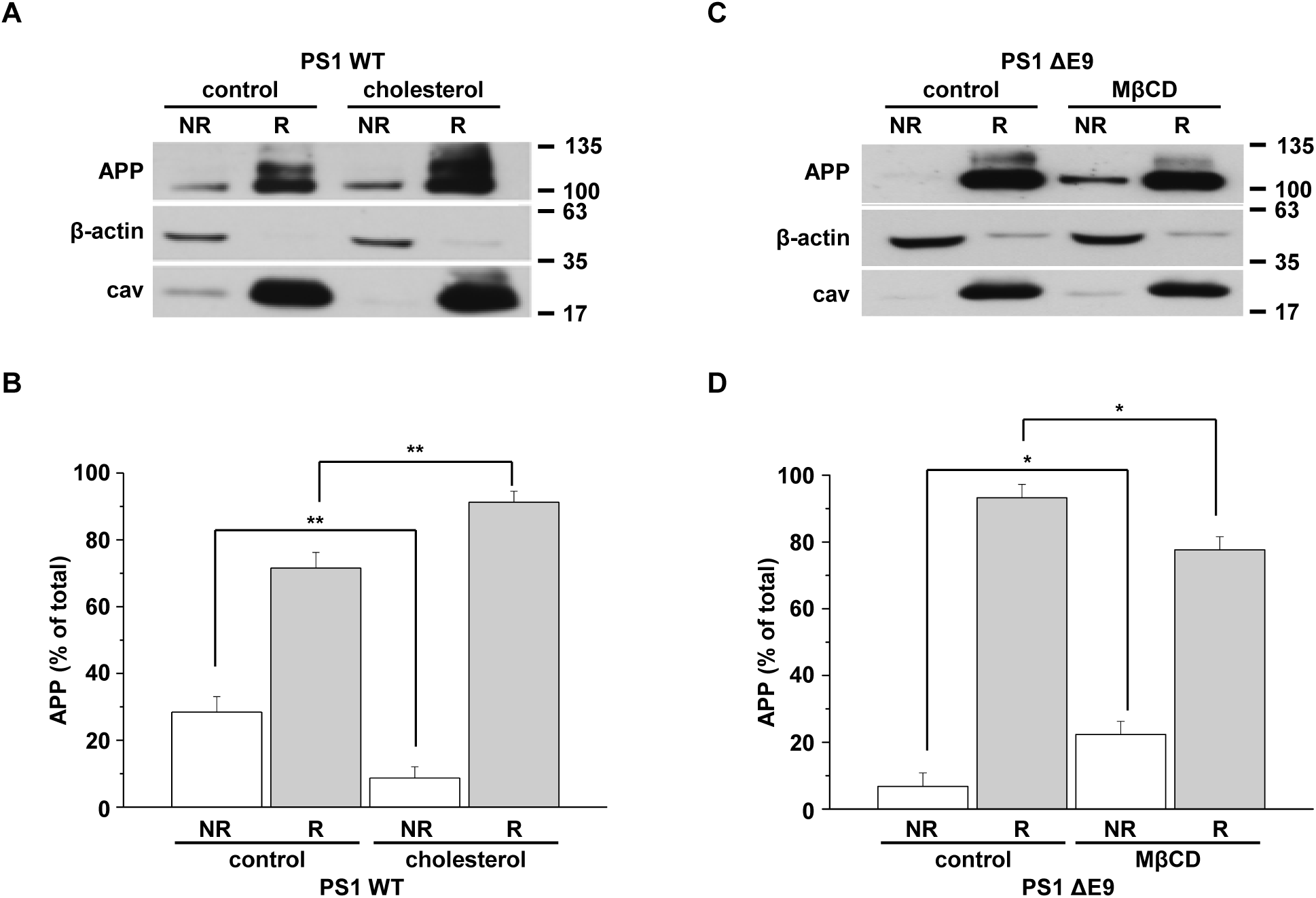
Cellular cholesterol levels determined the localization of surface APP in lipid raft microdomains. To examine APP localization at the plasma membrane, cells were incubated with EZ-Link NHS-biotin at 4°C to label all surface proteins. Then, biotin-labeled cell lysates were pooled for discontinuous sucrose density gradient to separate lipid raft and non-lipid raft fractions. Fractions 4 to 6 (lipid rafts, R) or 8 to 12 (non-lipid rafts, NR) were combined, and the same amount of protein was used for capturing biotin-labeled proteins with streptavidin beads. The biotin labeled proteins were run on western blot, and detected with APP, β-actin, and caveolin (lipid raft marker) antibodies. A. CHO PS1 WT cells were pre-treated with 150 μM MβCD-cholesterol before biotin-labeling. A typical result demonstrates surface APP levels from raft and non-raft fractions (n=6). B. Quantitative analysis of surface APP from PS1 WT control and cholesterol-treated PS1 WT cells (n=6). C. CHO PS1 ΔE9 cells were pre-treated with 5 mM MβCD to decrease cholesterol levels before labeling surface proteins. Representative western blot result indicates surface APP levels from raft and non-raft fractions (n=5). D. Quantitative analysis of surface APP from PS1 ΔE9 control and MβCD-treated PS1 ΔE9 cells (n=5). Statistical analysis was carried out by one-way ANOVA: *p<0.05, **p<0.01.

To investigate the effect of increasing cholesterol levels on surface APP localization in lipid raft fractions, CHO PS1 WT cells were pre-treated with 150 μM MβCD-cholesterol for 1 h (Fig 4A). APP levels in raft fractions were significantly increased to 91.3 ± 3.3% (n=6) by increasing cholesterol levels (Fig 4B). Conversely, surface APP localization in non-raft fractions was significantly decreased by MβCD-cholesterol. Next, CHO PS1 ΔE9 cells were pre-treated with 5 mM MβCD for 30 min to decrease cholesterol levels before labeling surface proteins. In PS1 ΔE9 control cells, most surface APP was localized in lipid raft fractions and barely detectable in non-lipid raft fractions (Fig 4C). Reducing cholesterol level increased the localization of surface APP in non-lipid raft fractions. The ratio of APP in non-lipid raft fractions increased from 6.8 ± 4.0% in control cells to 22.3 ± 3.9% (n=5) in cells treated with MβCD (Fig 4D). In contrast, the ratio of APP in lipid raft fractions showed a significant decrease from 93.2 ± 4.0% to 77.7 ± 3.9% (n=5) by MβCD treatment. These results indicated that modulating cholesterol levels regulates surface APP localization in lipid rafts, which could affect the further processing of APP and subsequent Aβ42 generation.

### 6. APP preferentially internalizes from lipid rafts

Our findings showed that cholesterol levels regulated the localization of surface APP in lipid rafts as well as the rate of APP endocytosis. However, it is still unclear whether lipid raft microdomains are the sites where APP undergoes endocytosis. To investigate this possibility, we applied a reversible biotinylation method to the internalization assay. Cells were pre-treated with EZ-Link NHS-SS-biotin to label surface proteins, and then further incubated at 37°C for 10 min to allow internalization of biotin-labeled surface proteins as described in Methods. All remaining surface biotins were then removed using reducing agents. Equal amount of biotin-bound proteins from cell lysates were loaded on discontinuous sucrose density gradient to obtain 12 fractions. The equal amount of proteins from lipid raft fractions (R; fractions 4 to 6), and non-raft fractions (NR; fractions 8 to 12) were captured with streptavidin beads to pull down biotin-bound proteins. Then, equal volumes of lipid raft and non-lipid raft fractions were loaded for Western blots. If the characteristics of the internalized membranes from lipid raft and non-raft fractions were maintained, the bead-captured APP represented the internalized APP originating from either lipid rafts or non-rafts during the 10 min incubation time. A typical result is shown in Fig 5A and 5C. The ratios of internalized APP from lipid raft and non-raft fractions were calculated and shown in Fig 5B and 5D. The ratio of APP internalized from lipid raft fractions was higher in PS1 ΔE9 cells (79.3 ± 2.1%, n=6) compared to PS1 WT cells (65.5 ± 4.7%, n=5). Considering that surface APP level in raft fractions was higher in PS1 ΔE9 cells than in PS1 WT cells (Fig 4), this result may suggest that APP is preferentially internalized from lipid rafts.

**Figure 5.**
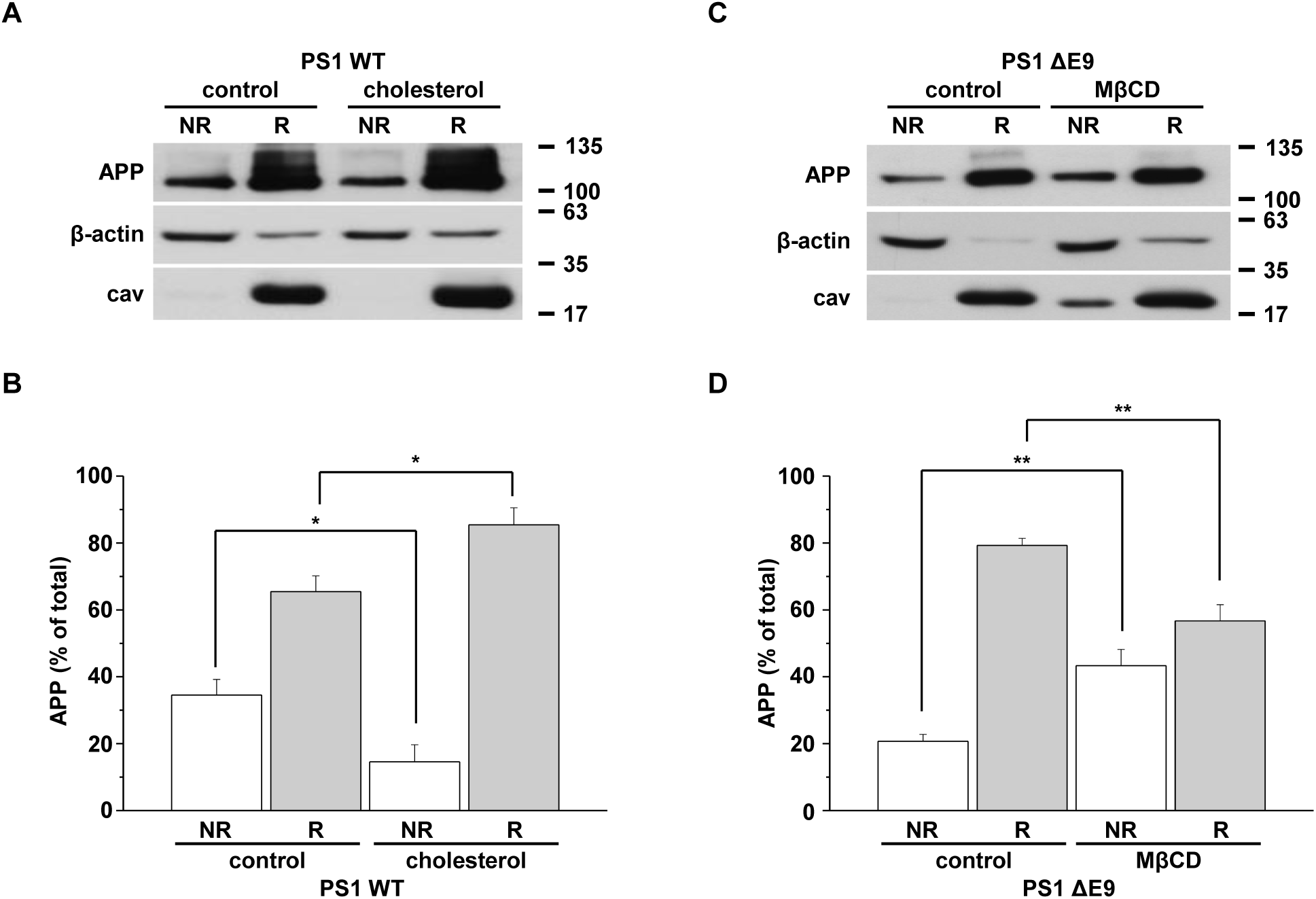
Cellular cholesterol levels determined raft-dependent endocytosis of APP. To verify the contribution of lipid rafts to APP endocytosis, cells were incubated with EZ-Link NHS-SS-biotin at 4°C to label surface proteins. Then, cells were incubated at 37°C for 10 min to allow internalization of biotin-labeled surface proteins. After internalization, remaining surface-bound biotin was removed, and biotin-labeled cell lysates were loaded for discontinuous sucrose density gradient centrifugation to separate lipid raft and non-lipid raft fractions. Fractions 4-6 (lipid rafts, R) or 8-12 (non-lipid rafts, NR) were pooled, and biotin-labeled proteins were captured with streptavidin beads. Internalized biotin-labeled proteins were loaded on western blots, and detected with APP, β-actin, and caveolin (lipid raft marker) antibodies. A. CHO PS1 WT cells were pre-treated with 150 μM MβCD-cholesterol before labeling surface proteins. Internalized biotin-bound APP levels from raft and non-raft fractions were detected by western blotting (n=5). B. Analysis of band densitometry shows the levels of internalized labeled APP from PS1 WT control and cholesterol-treated PS1 WT cells (n=5). C. CHO PS1 ΔE9 cells were pre-treated with 5 mM MβCD to reduce cholesterol levels before biotin-labeling. Internalized labeled APP from lipid raft and non-lipid raft fractions were monitored by western blot (n=6). D. The band densitometry of internalized labeled APP from lipid raft and non-lipid raft fractions was analyzed from PS1 ΔE9 and MβCD-treated PS1 ΔE9 cells (n=6). Statistical analysis was carried out by one-way ANOVA: *p<0.05, **p<0.01.

To investigate the effects of cholesterol level on APP internalization from lipid raft fractions, PS1 WT cells were pre-treated with 150 μM MβCD-cholesterol for 1 h. A typical result is shown in Fig 5A, where the ratio of internalized APP from lipid raft fractions was increased to 85.4 ± 5.1% (n=5) by MβCD-cholesterol. In contrast, when we decreased cholesterol levels from PS1 ΔE9 cells, the ratio of internalized APP from raft fractions was decreased to 56.7 ± 4.9% (n=6). Thus, increasing or decreasing localization of surface APP in lipid rafts consistently induced the same changes in the ratio of internalized APP. These results suggested that cholesterol level affected APP endocytosis from lipid rafts by regulating APP localization in these specific microdomains.

To test whether these cholesterol effects on APP endocytosis were cell-type dependent, we used HeLa cells stably transfected with APP751 carrying the Swedish mutation (APPswe). Firstly, cells were incubated with 150 μM MβCD-cholesterol for 1 h. Then, the biotin-labeled lipid raft fractions were obtained to monitor the membrane localization of surface APP. As shown Appendix Figure S3, increasing cellular cholesterol levels significantly increased the ratio of surface APP in lipid raft microdomains from 77.3 ± 3.4% to 90.5 ± 2.5% (n=5). The effect of cholesterol on lipid raft-dependent APP endocytosis was also tested. The ratio of APP internalized from lipid rafts was increased by elevating cholesterol levels (Appendix Figure S4), which was consistent with results from CHO PS1 WT cells. Cells were also pre-treated with 1 mM MβCD for 30 min to decrease cholesterol levels before labeling surface proteins using EZ-Link NHS-SS-biotin. Lowering cellular cholesterol levels significantly decreased the ratio of surface APP in lipid rafts from 81.9 ± 1.9% to 72.2 ± 3.0% and increased the ratio of surface APP in non-lipid rafts from 18.1 ± 1.9% to 27.8 ± 3.0% (n=5) (Appendix Figure S3). Internalized APP from raft fractions was significantly reduced by MβCD (Appendix Figure S4). From these results, we confirmed that the effects of cholesterol on APP localization in lipid raft fractions as well as on APP endocytosis from lipid rafts were not likely to be cell-type dependent.

### 7. Cholesterol increases accumulation of internalized APP in early endosomes

Newly synthesized and matured APP from endoplasmic reticulum (ER) and trans-Golgi network (TGN) is transported to plasma membrane (Thinakaran & Koo, 2008). At the plasma membrane, APP is either cleaved by α-secretase, or undergoes endocytosis through clathrin-dependent and -independent endocytosis within minutes. Since we confirmed that APP endocytosis was sensitive to cellular cholesterol levels, we next determined its subcellular localization. To achieve this, surface APP was labeled with 6E10 antibody in CHO PS1 WT or PS1 ΔE9 cells before internalization. After allowing 5 or 10 min for the internalization of labeled APP at 37°C, cells were fixed and remaining surface APP was captured by anti-mouse secondary antibody to eliminate surface signals. Next, cells were permeabilized, and incubated with EEA1 (early endosome marker) antibody for 2 h. After washing, internalized APP and early endosomes were labeled with Alexa647-, and Alexa488-conjugated secondary antibodies, respectively. Typical fluorescence reactivities for APP (red) and EEA1 (green) are shown in Fig 6A. The coefficient of internalized APP and EEA1 was analyzed in Fig 6B. The co-localization of APP and EEA1 was much higher in PS1 ΔE9 cells compared to PS1 WT cells both at 5 min and 10 min (n=4), showing the increased APP localization in early endosomes. However, the slopes of coefficient increase from 5 min to 10 min were almost identical in both cells, indicating that the rate of APP trafficking from the plasma membrane to early endosomes was similar in both cells. Thus, our results suggest that only the amount of internalized APP was increased in PS1 ΔE9 cells compared to PS1 WT cells.

**Figure 6.**
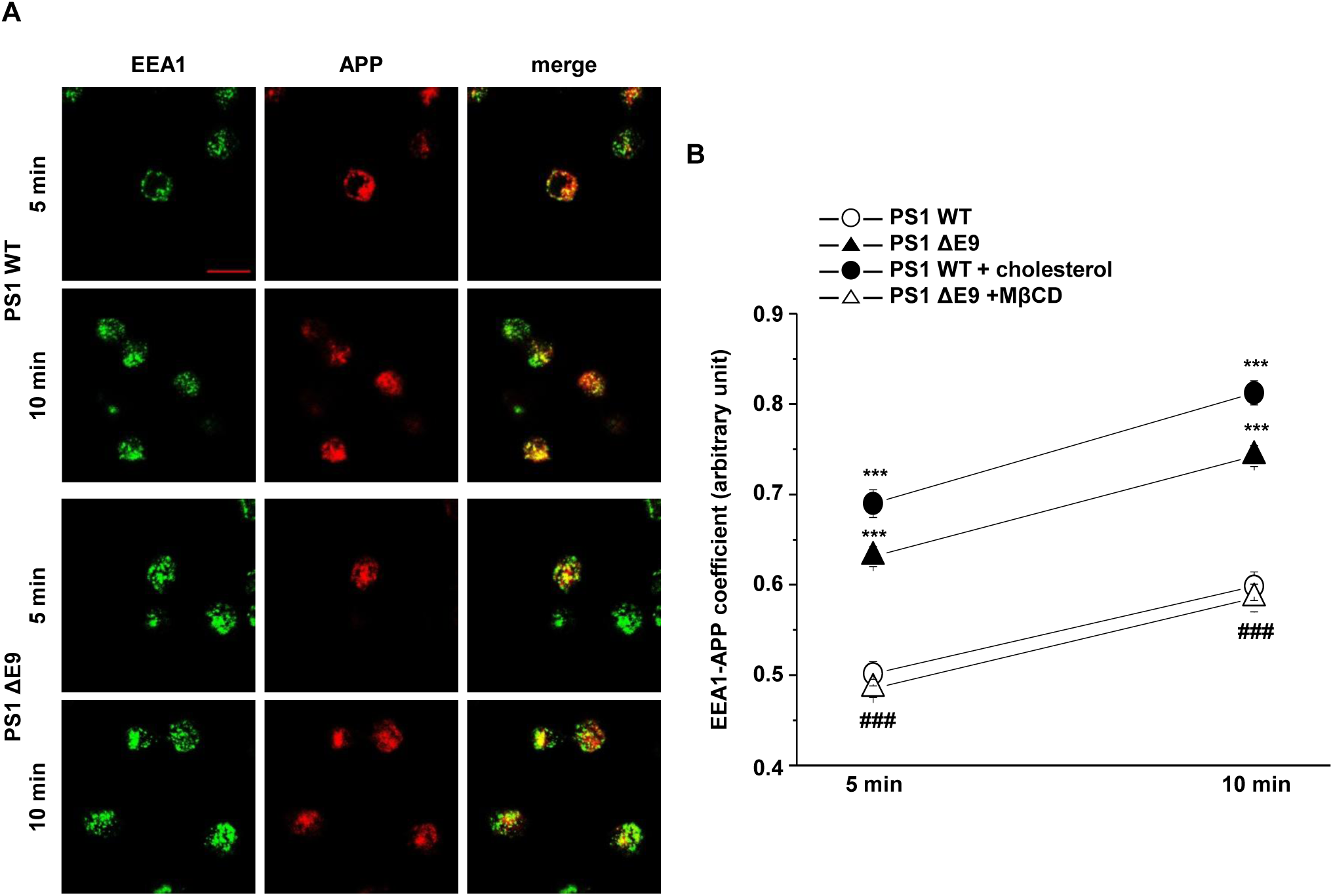
Cellular cholesterol levels altered the accumulation of APP in early endosomes. CHO PS1 WT and PS1 ΔE9 cells were incubated with APP antibody at 4°C and transferred to 37°C for indicated times to allow internalization of the labeled surface APP. Then, cells were fixed and surface APP was stained with anti-mouse IgG secondary antibody to eliminate remaining surface APP signal. Following permeabilization, cells were labeled with EEA1 antibody to label early endosomes. Subsequently, anti-mouse Alexa647 (red)- or anti-rabbit Aexa488 (green)-conjugated secondary antibodies were used to label internalized APP and early endosomes, respectively. A. Representative confocal image showing APP localization at each time. Data are representative of four independent experiments. Scare bars correspond to 10 μm. B. Fluorescence intensities of APP and EEA1 were measured using Image J. The co-efficiency of APP and early endosomes was determined with Image J (n=4). Statistical analysis was analyzed by one-way ANOVA: *** indicates P<0.001 significant difference compared to PS1 WT cells. ### represents P<0.001 significant difference compared to PS1 ΔE9 cell.

We also investigated the effects of regulating cholesterol levels on subcellular localization of APP. PS1 WT cells were pre-treated with 150 μM MβCD-cholesterol for 1 h, and internalized APP was monitored. Typical fluorescence reactivities for APP and EEA1 are shown in Fig EV4. The co-localization of APP and EEA1 is shown in Fig 6B. MβCD-cholesterol significantly increased the amount of APP accumulated in early endosomes by 1.4-fold without change in the slope of coefficient from 5 min to 10 min (n=4). In this experiment, APP-EEA1 coefficient looked very similar to that of PS1 ΔE9 cells. We performed the reverse experiment by reducing cholesterol levels. PS1 ΔE9 cells were pre-treated with 5 mM MβCD for 30 min, and internalized APP was monitored. Typical fluorescence reactivities for APP and EEA1 are shown in Fig EV4. Reducing cholesterol levels significantly decreased the amount of APP accumulated in early endosomes (Fig 6B; n=4). Again, APP-EEA1 coefficient looked very similar to that of PS1 WT cells. These findings indicate that cholesterol levels were closely associated with the amount of internalized APP reaching early endosomes from the plasma membrane.

### 8. Localization of endogenous neuronal APP in lipid raft microdomains at the plasma membrane is determined by cellular cholesterol levels

To test whether cholesterol levels in neurons affected the partitioning of surface APP into lipid raft microdomains, we used primary hippocampal neurons from Sprague-Dawley rat embryos. Previous research showed that these neurons are fully matured at 21-35 days in vitro (DIV21-DIV35; stationary phase), and are characterized by pyramidal shaped cell bodies and intensive connections and networking of neurites (Bertrand et al., 2011). Neurons (DIV21-DIV23) were incubated with 2 mM MβCD or 1.5 mM MβCD-cholesterol for 30 min to decrease or to increase cellular cholesterol levels, respectively. Then, neurons were incubated with 300 μg/ml filipin for 2 h to stain free cholesterol as shown in Fig 7A. Filipin fluorescent intensities were compared in Fig 7B. MβCD-cholesterol treatment elevated cholesterol levels by 33.3 ± 3.0% (n=3) while MβCD treatment decreased cholesterol levels by 21.2 ± 2.1% (n=3). To monitor the localization of APP at the plasma membrane, cells were co-stained with APP antibody and CTB, a lipid raft marker. After fixing and permeabilizing, neurons were immunostained with NeuN antibody for 2 h to identify neurons. Surface membrane of both cell bodies and axons which were covered with cholesterol-enriched myelin sheet were stained with CTB (Fig 7C, n=4). The endogenous neuronal APP was mostly localized in the cell bodies. Elevated cholesterol increased the co-localization of APP and CTB (Fig 7D, n=4). In contrast, MβCD decreased the co-localization of APP-CTB. These results suggested that lipid raft localization of APP was regulated by cholesterol levels, a finding consistent with the results obtained from APP-transfected cells.

**Figure 7.**
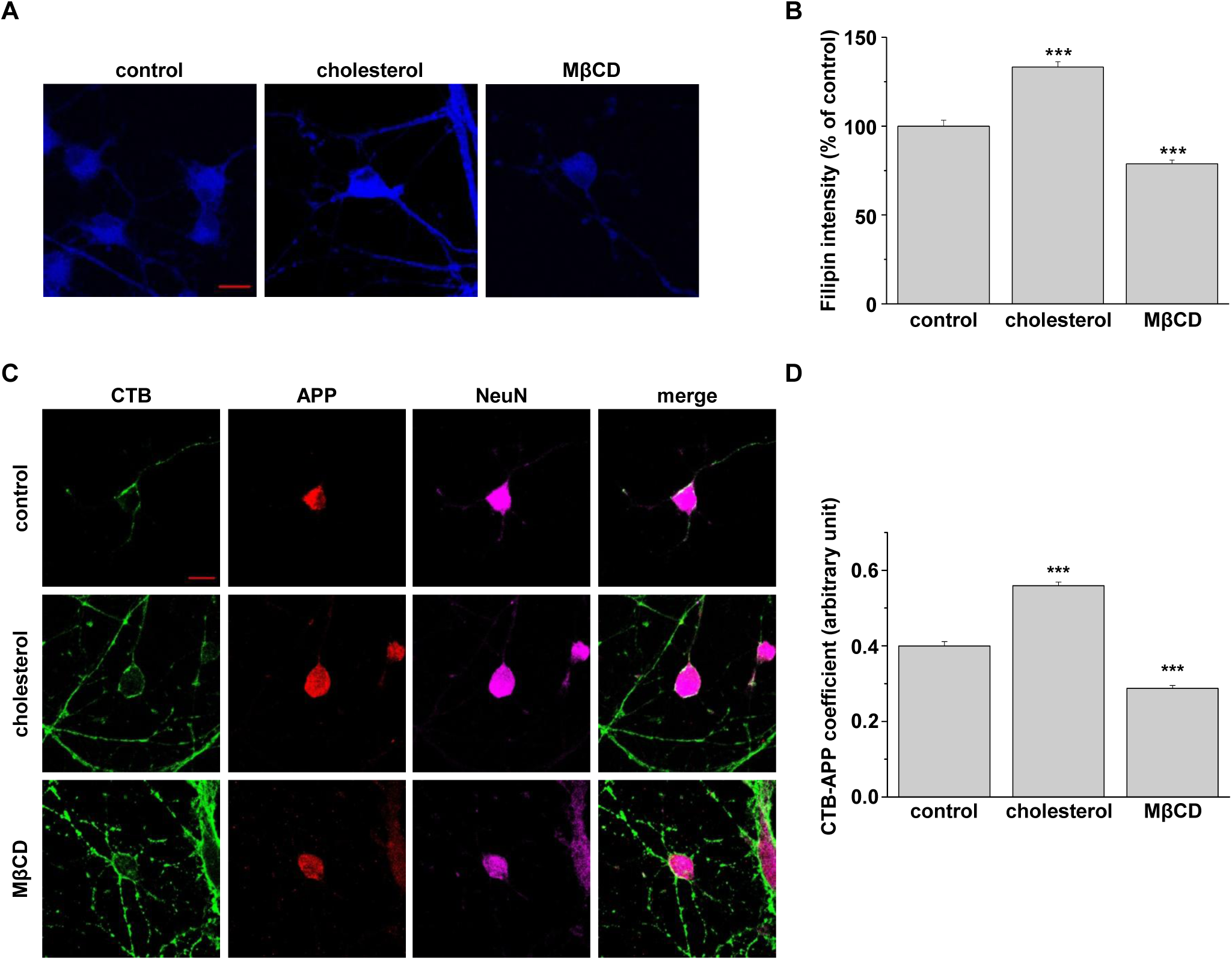
Cellular cholesterol levels determined the localization of endogenous APP in lipid raft microdomains from rat primary hippocampal neurons. A. Rat primary hippocampal neurons (DIV21-DIV23) were incubated with either 1.5 mM MβCD-cholesterol or 2 mM MβCD. Then, cells were incubated with filipin to stain free cholesterol levels. Representative confocal image are shown. Data are representative of three independent experiments. Scare bars correspond to 10 μm. B. Filipin fluorescent intensities (n=3). C. Hippocampal neurons were treated with MβCD-cholesterol or MβCD, followed by incubation with APP antibody and cholera toxin B (CTB) at 4°C. After fixing and permeabilization, neurons were incubated with NeuN antibody to detect neurons. Confocal image from four independent experiments is shown. Scare bars correspond to 10 μm. D. Co-efficiency of APP and CTB was analyzed with Image J (n=4). Statistical analysis was performed by one-way ANOVA: ***p<0.001.

## Discussion

Our data demonstrated that cholesterol recruited surface APP into lipid raft microdomains. Our findings also provided evidence of a direct relationship between APP localization in cholesterol-enriched lipid rafts and APP endocytosis as shown in Fig. 8 for our current model. Since Aβ is produced from internalized APP in early endosomes and lysosomes, regulating APP localization in lipid rafts could be a promising new therapeutic target for Alzheimer’s disease.

**Figure 8.**
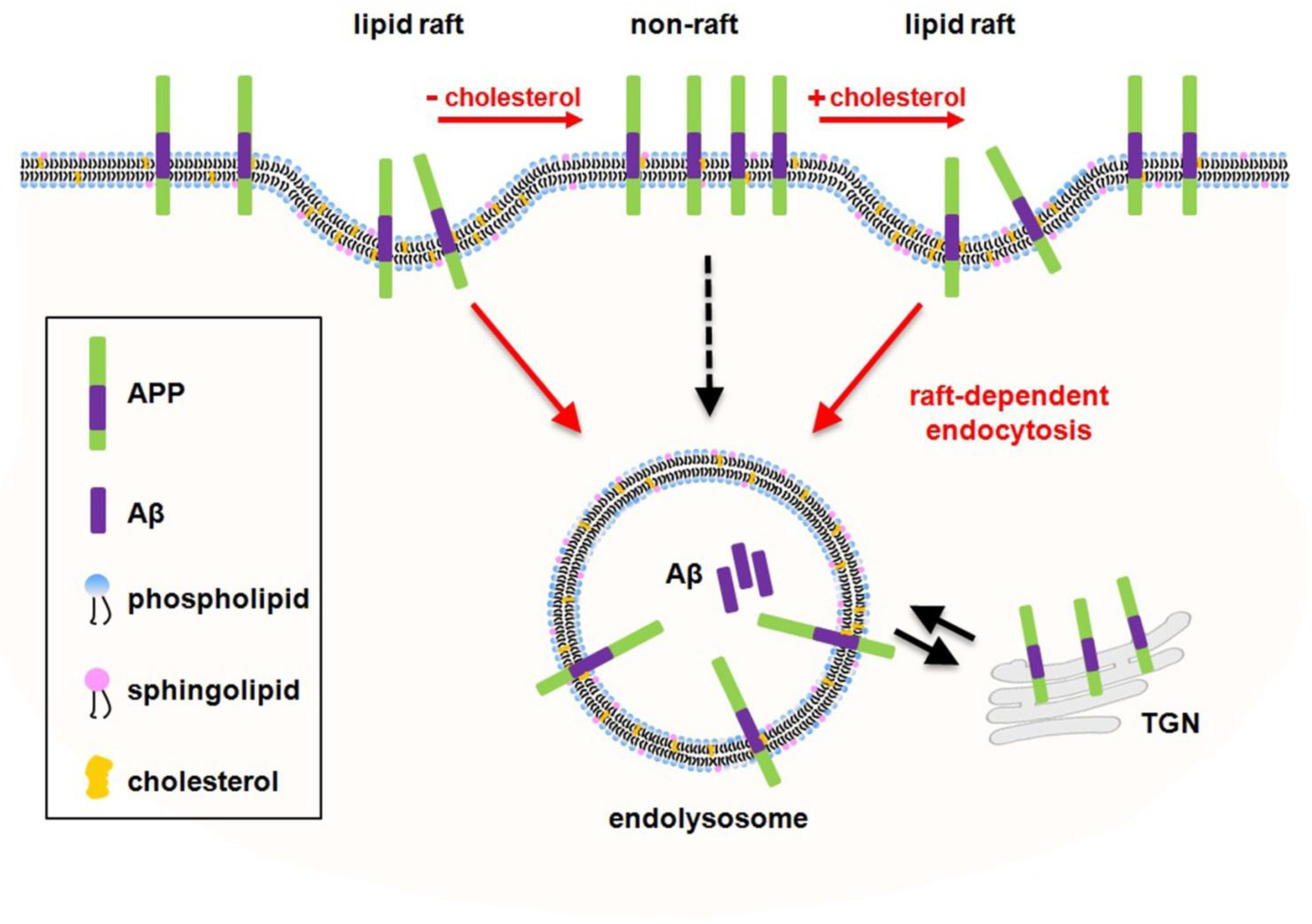
Current model for our study.

Many lines of evidence shed light on the role of cholesterol in AD (Allinquant et al., 2014, Maulik et al., 2013). Changes in cholesterol homeostasis as well as other lipid classes in postmortem AD brain are considered to represent a third pathological feature of AD (Fabelo et al., 2014, Heverin et al., 2004, Jarvik et al., 1995, Vanmierlo et al., 2010, Xiong et al., 2008). APP processing is highly influenced by membrane cholesterol since APP and the associated secretases are localized in cholesterol-enriched lipid raft structures (Benjannet et al., 2001, Bhattacharyya et al., 2013, Hicks et al., 2012, Hur et al., 2008, Osenkowski et al., 2008). Lipid rafts have also been implicated in the regulation of several membrane trafficking and cell signaling pathways (Kirkham et al., 2005, Kirkham & Parton, 2005a, Kirkham & Parton, 2005b, Okamoto et al., 2000). Lowering membrane cholesterol attenuates APP endocytosis and promotes non-amyloidogenic processing, while increasing membrane cholesterol levels escalates APP internalization into acidic intracellular compartments, enhancing Aβ production (Cossec et al., 2010, Kojro et al., 2001). These findings suggested that the cholesterol composition of the plasma membrane and the distribution of APP in lipid rafts are critical for APP processing and Aβ generation. Indeed, we found that increasing or decreasing cholesterol levels regulated Aβ42 production, consistent with changes induced in APP endocytosis and localization in lipid rafts (Fig EV5). When we increased cellular cholesterol levels in CHO PS1 WT cells, the level of secreted Aβ42 was significantly increased. In contrast, the increased Aβ42 levels seen in the presence of elevated cholesterol in CHO PS1 ΔE9 cells was significantly reduced by decreasing cellular cholesterol level.

Based on the co-localization evidence, elevated cholesterol levels in PS1 ΔE9 cells increased APP localization in both caveolin-and CTB-positive regions, which represent lipid raft microdomains. Manipulation of cellular cholesterol with MβCD or MβCD-cholesterol decreased or increased the degree of colocalization of APP-caveolin and APP-CTB, respectively. In addition to cell models expressing APP, we also used primary hippocampal neurons to show the effects of manipulating cellular cholesterol on the localization of endogenous APP in lipid raft microdomains. These findings demonstrated that surface APP localization within lipid rafts was regulated by cellular cholesterol levels. This result was consistent with our previous report in which elevated cellular cholesterol increased APP localization in lipid rafts (Cho et al., 2019). In addition, this lateral accumulation of APP in lipid rafts by the elevated cholesterol supports previous reports showing that cholesterol binds to the transmembrane region of APP/C99 and favors the formation of APP/C99-cholesterol complex (Barrett et al., 2012, Beel et al., 2008, Beel et al., 2010, Song et al., 2013, Song et al., 2014).

Endocytosis is a serial process of membrane dynamics. Therefore, membrane lipid composition is critical for endocytic mechanisms (Doherty & McMahon, 2009, El-Sayed & Harashima, 2013, Yue & Xu, 2015). Also, the distribution of membrane proteins between lipid raft and non-lipid raft microdomains is thought to regulate the endocytosis of certain proteins (Doherty & McMahon, 2009, El-Sayed & Harashima, 2013). Consistent with these results, we found that endogenously elevated cholesterol in CHO PS1 ΔE9 cells resulted in increased APP endocytosis compared to PS1 WT cells. However, the endocytosis of transferrin was not changed by the altered cholesterol levels. These results may suggest that general clathrin-dependent endocytosis is unaffected by the elevation in membrane cholesterol levels, while conversely, it did appear to be specific for the internalization of APP. The internalized APP was derived from both lipid raft and non-lipid raft microdomains. However, the elevated cholesterol level in CHO PS1 ΔE9 cells induced preferential internalization of APP from lipid rafts. When we depleted cholesterol levels with MβCD, the portion of internalized APP derived from lipid rafts was decreased. In addition, these cholesterol effects on APP localization and APP endocytosis were not cell-type specific since we recapitulated these effects using HeLa APPswe cells. We also found that modification of cholesterol content affected the amount of internalized APP reaching early endosomes. However, the rate of internalization was not affected by changing cholesterol levels. Thus, APP endocytosis itself, but not subsequent trafficking to early endosomes was altered by cholesterol levels.

According to Cossec and colleagues, APP endocytosis is increased by cholesterol in a clathrin-dependent manner (Cossec et al., 2010). Interestingly, our data showed that internalized APP originated from both raft and non-raft domains. In addition, APP internalization from both microdomains was sensitive to cholesterol modulation. It has been proposed that the lipid raft-associated protein, flotillin, can cluster with APP into lipid raft microdomains by binding to its C-terminal region (Chen et al., 2006, Schneider et al., 2008). APP can also bind cholesterol through its transmembrane domain (Barrett et al., 2012, Beel et al., 2008, Beel et al., 2010, Song et al., 2013, Song et al., 2014). Taken together, it is possible that APP associated with either cholesterol or flotillins, is recruited to lipid raft microdomains, and that recruited APP in lipid rafts may be internalized to the endosomal/lysosomal subcellular trafficking system. This form of APP endocytosis may occur via raft-dependent pathways such as caveolin- or flotillin-mediated endocytosis (Kirkham et al., 2005, Kirkham & Parton, 2005a, Kirkham & Parton, 2005b, Okamoto et al., 2000). Clathrin-dependent APP endocytosis is also affected by cellular cholesterol (Cossec et al., 2010, McMahon & Boucrot, 2011, Subtil et al., 1999), which is consistent with our result showing that the internalized APP from non-rafts was also changed by alterations of cholesterol levels. Considering the previous reports, our results may suggest that APP can be internalized through multiple pathways including both classical clathrin-dependent endocytosis and raft-induced endocytosis. Further studies would be needed to test this possibility.

## Materials & Methods

### Cell culture and experimental treatments

Wild-type human APP751 expressing Chinese hamster ovary (CHO) cells (Kang et al., 2013) were stably transfected with either presenilin 1 wild type (PS1 WT) or ΔE9 mutant (PS1 ΔE9). Stable CHO PS1 WT and PS1 ΔE9 cell lines were grown in Dulbecco’s Modified Eagle Medium (DMEM) supplemented with 10% (v/v) heat-inactivated fetal bovine serum (FBS), 100 U/ml penicillin, 100 μg/ml streptomycin, and 250 μg/ml Zeocin at 37°C with 5% CO_2_ atmosphere. For cholesterol modulation, CHO PS1 WT cells were incubated with 150 μM MβCD-cholesterol (cholesterol-water soluble; Sigma, #C4951) for 1 h and CHO PS1 ΔE9 cells were treated with 5 mM methyl-β-cyclodextrin (MβCD; Sigma, #332615) for 30 min.

HeLa cells stably expressing APP751 carrying the Swedish mutation (APPswe) were maintained in DMEM with 10% heat-inactivated FBS, 100 units/ml penicillin, 100 μg/ml streptomycin, 250 μg/ml Zeocin, and 400 μg/ml G418 at 37°C with 5% CO_2_ atmosphere. HeLa APPswe cells were incubated with either 1 mM MβCD or 150 μM MβCD-cholesterol for 30 min to reduce or increase cellular cholesterol levels, respectively.

### Rat primary hippocampal neuron culture

Embryonic 18-day-old Sprague-Dawley rat fetuses were prepared for rat primary hippocampal neurons. All procedures were carried out in accordance with the guidelines of Sungkyunkwan University Animal Care and Ethics Committee. Rat primary hippocampus from both hemispheres were dissected and rinsed with Hank’s balanced salt solution (HBSS). Then, hippocampi were incubated with HBSS solution containing 0.25% trypsin-EDTA at 37°C for 5 min. Next, the tissue was incubated with 1 ml FBS in 4 ml HBSS at 37°C for 3 min to inactivate trypsin-EDTA and then washed three times with HBSS. Next, tissue was resuspended in neurobasal medium and filtered through 100 μm strainer to dissect tissue into single neurons. Then, neurons were washed with HBSS and resuspended in neurobasal medium supplemented with 2% B27, 2 mM glutamax, and 1% penicillin/streptomycin (all supplements from GIBCO). Finally, primary hippocampal neurons were grown on poly-D-lysine coated glass cover slips, and neurons were maintained for 21 to 23 days in vitro (DIV21-DIV23). For cholesterol manipulation, neurons were treated with either 1.5 mM MβCD-cholesterol or 2 mM MβCD for 30 min to increase or decrease cholesterol, respectively.

### Co-localization experiments

Cells were grown on poly-D-lysine coated cover glass. After washing with ice-cold phosphate-buffered saline (PBS), cells were incubated with APP antibody (6E10; BioLegend, monoclonal, #803002) at 4°C for 1 h to label cell surface APP. Then, cells were fixed with 4% paraformaldehyde in 4% sucrose for 15 min and permeabilized in PBS containing 0.1% Triton X-100 and 2% bovine serum albumin (BSA) for 5 min. After washing, cells were blocked with 1% BSA in PBS solution for 1 h, and then cells were incubated with caveolin antibody (cav-1; BD Transduction Laboratories, polyclonal, #610059) for 2 h in blocking buffer to detect lipid raft microdomains. Following washing with PBS, cells were incubated with goat anti-mouse conjugated with Alexa Fluor 488 (Invitrogen, #A11001) and donkey anti-rabbit conjugated with Alexa Fluor 647 (Invitrogen, #A31573) secondary antibodies in blocking buffer at 4°C for 16 h to label primary antibodies APP and cav-1, respectively. Next day, cells were washed with PBS and mounted with mounting medium (Sigma, #F6182). Immunofluorescence staining was monitored on a confocal microscope (LSM710, Zeiss). For co-localization with cholera toxin B (CTB), CHO cells were incubated with 6E10 and 10 μg/ml FITC-conjugated cholera toxin B subunits (Sigma, #C1655) to label lipid raft microdomains at 4°C for 1 h before fixation. Then, cells were processed following steps as described above. APP was captured with goat anti-mouse conjugated with Alexa 647 (Invitrogen, #A21240) secondary antibody.

Primary hippocampal neurons were grown on poly-D-lysine cover slips and stained with 6E10 to label endogenous APP at the plasma membrane and 7 μg/ml FITC-conjugated CTB subunits to label lipid raft microdomains at 4°C for 1 h. Then, neurons were fixed with 4% paraformaldehyde in 4% sucrose for 15 min and permeabilized in PBS containing 0.1% Triton X-100/2% BSA for 5 min. After 1 h blocking with 2% BSA in PBS, neurons were incubated with α-NeuN antibody (Millipore, polyclonal, #ABN78) for 2 h at room temperature to detect neurons. Following washes with PBS, cells were incubated with goat anti-mouse conjugated with Alexa Fluor 568 (Invitrogen, #A11004) and donkey anti-rabbit conjugated with Alexa Fluor 647 in blocking buffer overnight to detect primary antibodies 6E10 and α-NeuN, respectively. Next day, cells were washed and mounted with mounting medium. Immunofluorescence reactivity was captured on a confocal microscope (LSM710, Zeiss). The co-localization of APP and caveolin or APP and CTB was calculated using the JACop plug-in of Image J program (https://imagej.net/Colocalization_Analysis), which is based on the Manders correlation coefficient (Dunn, Kamocka et al., 2011).

### Primary antibody uptake assay

CHO PS1 WT and PS1 ΔE9 cells were grown on glass cover slips coated with poly-D-lysine. The next day, the cells were washed three times with ice-cold PBS for 5 min, and were incubated with 6E10 antibody (1:100) in PBS containing 2% BSA for 1 h at 4°C to label surface APP at the plasma membrane. Next, cells were washed with ice-cold PBS, and cells were transferred to 37°C for various times to allow internalization. At 0 min, cells were fixed with 4% paraformaldehyde in 4% sucrose for 15 min, followed by incubation with goat anti-mouse Alexa647 (red)-conjugated secondary antibody at 4°C for 1 h to label surface APP. After internalization, cells were fixed and incubated with goat anti-mouse Alexa647 (red)-conjugated secondary antibody at 4°C for 1 h to label surface APP. Next, cells were washed with PBS and then permeabilized in PBS containing 0.1% Triton X-100 and 2% BSA for 5 min. Cells were blocked with 1% BSA in PBS for 1 h at room temperature. Cells were washed with PBS twice, and incubated with goat anti-mouse Alexa488 (green)-conjugated secondary antibody in blocking buffer for 16 h to label internalized APP. The following day, cells were washed with PBS and mounted with mounting medium and left overnight at 4°C to dry. Immunofluorescence reactivity was captured with a confocal microscope (LSM710, Zeiss). Mean fluorescence intensity values corresponding to plasma membrane as measured from the edge of the cell to a depth of 500 nm (surface APP; A) and mean fluorescence intensity values from the region of interest (ROI) according to the cell shape (internalized APP; B) were measured using the Image J program. The rate of APP endocytosis was obtained by calculating the ratio of B/A.

### Transferrin uptake

Cells were incubated with 25 μg/ml Alexa Fluor 488-conjugated transferrin (Invitrogen, #T-13342) in PBS containing 0.1% BSA for 5, 10, 30 min at 37°C. Surface transferrin was removed from the plasma membrane by treating cells with pH 5.5 buffer (0.1 M sodium acetate, 0.05 M NaCl) for 5 min. Cells were then washed with PBS and fixed with 4% paraformaldehyde in 4% sucrose for 15 min at room temperature. Immunofluorescence reactivity was captured using a confocal microscope (LSM710, Zeiss). The Image J program was used to measure mean fluorescence intensity values from the region of interest (ROI) according to the cell shape.

To quantitatively measure fluorescence intensity of transferrin internalization, cells were incubated in PBS with 0.1% BSA containing 25 μg/ml Alexa Fluor 488-conjugated transferrin for 5, 10, 30 min at 37°C. Then, cells were washed with PBS and lysed with lysis buffer (50 mM Hepes. pH 7.2, 100 mM NaCl, 1% Triton X-100, 1 mM sodium orthovanadate, and protease inhibitor mixture). The same amount of cell lysate protein was loaded on Tris-glycine SDS-PAGE gel. Proteins were transferred to nitrocellulose membrane, and the intensity of the bands was detected using LAS 3000 (Fuji Film). Band densitometry was analyzed using Multi Gauge V3.0 program.

### Reversible biotinylation assay

CHO cells were washed with ice-cold PBS and incubated in PBS supplemented with 0.25 mg/ml sulfo-NHS-SS-biotin (Thermo, #21441) for 10 min at 4°C. Excess biotin was washed out with ice-cold PBS containing 100 mM glycine. Cells were incubated with 1% BSA in PBS for 15 min at 4°C. After washing with PBS, cells were incubated at 37°C for appropriate times. For the zero minute time point, cells were kept at 4°C as a control. Cells were quickly washed in ice-cold PBS to stop internalization. Remaining cell surface biotin was cleaved off by incubating twice with reducing agent (50 mM sodium-2-mercapoethanesulfomate, 150 mM NaCl, 1 mM EDTA, 0.2% BSA, 20 mM Tris HCl, pH 8.6) for 25 min at 4°C. This reaction was quenched by ice-cold PBS containing 5 mg/ml iodoacetamide (Sigma, #I1149) with 1% BSA for 10 min. We also incubated cells in PBS supplemented with 0.25 mg/ml sulfo-NHS-biotin (Thermo, #21217) at 4°C to detect all surface proteins. For total pool of surface APP, this sample does not undergo the above reducing and quenching steps. After washing, cells were extracted in lysis buffer (50 mM Hepes, pH 7.2, 100 mM NaCl, 1% Triton X-100, 1 mM sodium orthovanadate, and protease inhibitor mixture). The equal amount of proteins were incubated with streptavidin-agarose slurry (Millipore, #16-126) at 4°C for 16 h in order to pull down all biotin-labeled proteins. After washing, the bound material was analyzed by Western blot. We calculated the rate of APP internalization as described below.

A: the levels of total surface APP (sulfo-NHS-biotin incubated, non-reduced and non-quenched)

B: control (0 min; sulfo-NHS-SS-biotin incubated, kept at 4°C)

C: internalized APP at 37°C (sulfo-NHS-SS-biotin incubated and internalized 5, 10, or 30 min) The (C-B)/A ratio represented the rate of internalized APP during each time point.

### Western blotting

The proteins were loaded on 8-10% Tris-glycine SDS-PAGE gel. Proteins were transferred to 0.2 μm nitrocellulose membrane and the transferred membrane was blocked with 5% (w/v) non-fat dried milk in Tris-buffered saline with 1% Tween-20 (TBST) for 1 h at room temperature. After washing blocked membrane with PBS four times for 10 min, the membrane was incubated with the following primary antibodies: APP (6E10; BioLegend, monoclonal, #803002), caveolin (cav-1; BD Transduction Laboratories, polyclonal, #610059), flotillin-1 (BD Transduction Laboratories, monoclonal, #610880), GAPDH (Cell Signaling Technology, monoclonal, #14C10), β-actin (EnoGene, monoclonal, #E12-041) at 4°C for 16 h. Next, the membrane was washed with TBST four times and incubated with horseradish peroxidase-conjugated goat anti-rabbit IgG (Invitrogen, polyclonal, #656120) or goat anti-mouse IgG (Invitrogen, polyclonal, #G21040) antibodies for 1 h at room temperature to detect each primary antibody. After the incubation with the secondary antibody, the membrane was washed again with TBST four times. We used enhanced chemiluminescence reagent (Westsave, #LF-QC0101), and signals were captured with film (MTC Bio, #A8815). The intensity of bands was captured by LAS-3000 system (Fuji Film) and analyzed by Multi Gauge V3.0.

### The localization of surface APP in lipid raft microdomains

To monitor the localization of cell surface APP between lipid raft and non-lipid raft microdomains, cells were incubated with PBS supplemented with 0.25 mg/ml sulfo-NHS-biotin for 10 min at 4°C to label all cell surface proteins. Remaining biotins were washed away with 100 mM glycine in PBS three times. Then, cells were collected with 0.25% trypsin-EDTA and lysed in 4-morpholineethanesulfonic acid (MES)-buffered saline (MBS; 25 mM MES, 150 mM NaCl, pH 6.5) containing 500 mM sodium carbonate (Sigma, #S7795) and a protease inhibitor cocktail Set III (Calbiochem, #535140). The lysates were homogenized 20 times with a 2 ml homogenizer and sonicated for 1 min (20 s sonication followed by 10 s interval). Cells were not homogenized with needle for this experiment. Equal amounts of protein were added to 0.8 ml of 80% (w/v) sucrose in MBS. Then, 1.6 ml of 35% (w/v) sucrose and 5% (w/v) sucrose in MBS were layered in a 5.1 ml ultracentrifuge tube (Beckman Coulter, #326819) to form a discontinuous sucrose gradient. The tubes were placed in a Beckman SW 55 Ti rotor (Beckman Coulter) and centrifuged at 50,000 rpm for 3 h at 4°C. From the top to the bottom, 12 fractions (0.4 ml each) were collected. Fractions #4-6 were combined as lipid raft fractions and fractions #8-12 were combined as non-lipid raft fractions. Equal amounts of protein from lipid raft and non-lipid raft fractions were incubated with streptavidin-agarose slurry at 4°C for 16 h to pull down biotin-labeled proteins. After washing, the biotin-labeled proteins were analyzed by Western blot to detect APP, β-actin, GAPDH, and lipid raft markers caveolin and flotillin.

### Rate of endocytosis of surface APP in lipid raft microdomains

To monitor the contribution of lipid raft microdomains to APP endocytosis, cells were washed with ice-cold PBS to block protein trafficking at the surface level. All procedures were performed on ice. Cells were incubated with PBS supplemented with 0.25 mg/ml sulfo-NHS-SS-biotin for 10 min at 4°C to label all surface proteins. Excess biotin was washed out, and cells were incubated with PBS containing 1% BSA for 15 min at 4°C. After washing, cells were incubated at 37°C for 10 min to allow internalization of biotin-labeled surface proteins. Then, cells were quickly placed on ice and washed with ice-cold PBS to stop internalization. Remaining cell surface biotin was removed by twice incubating cells with reducing agent (50 mM sodium-2-mercapoethanesulfomate, 150 mM NaCl, 1 mM EDTA, 0.2% BSA, 20 mM Tris HCl, pH 8.6) for 25 min at 4°C. Then, reducing agent was quenched by ice-cold 5 mg/ml iodoacetamide in 1% BSA for 10 min. After washing, cells were harvested with 0.25% trypsin-EDTA. Cells were lysed MBS containing 500 mM sodium carbonate with a protease inhibitor cocktail. The lysates were homogenized 20 times with 2 ml homogenizer followed by sonication for 1 min (20 s sonication followed by 10 s interval). Cells were not homogenized with needle for this experiment. After discontinuous sucrose gradient centrifugation, biotin-labeled internalized proteins were pulled down from fractions as described above. APP, β-actin, GAPDH, caveolin and flotillin were monitored by Western blot.

### Localization of APP in early endosomes

CHO PS1 WT and PS1 ΔE9 cells were grown on poly-D lysine coated glass cover slips. Surface APP was labeled with 6E10 antibody at 4°C for 1 h, and then, cells were transferred to 37°C in order to allow internalization. Internalization was stopped with ice-cold PBS and cells fixed with 4% paraformaldehyde in 4% sucrose at room temperature for 15 min. After washing, cells were incubated with goat anti-mouse IgG secondary antibody to captured remaining surface APP, which eliminated the surface signal. Internalized APP was captured after permeabilization. Next, cells were permeabilized with 0.1% Triton X-100/2% BSA/PBS buffer for 5 min. Then, cells were blocked with 2% BSA in PBS for 1 h. Cells were incubated with an antibody to early endosome marker, EEA1 (Cell Signaling, monoclonal, #C45B10) in blocking buffer for 2 h at room temperature. Following PBS washes, cells were incubated with goat anti-mouse conjugated with Alexa Fluor 647 and goat anti-rabbit conjugated with Alexa Fluor 488 (Invitrogen, #A1S1034) secondary antibodies in blocking buffer overnight in order to detect internalized APP and early endosomes, respectively. Next day, cells were washed with PBS and mounted with mounting medium. Immunofluorescence staining was monitored on a confocal microscope (LSM710, Zeiss). The co-localization of APP and early endosomes were measured by Image J program.

### Aβ42 peptide ELISA assay

CHO PS1 WT cells were pre-treated with 0, 75, or 150 μM MβCD-cholesterol for 1 h and CHO PS1 ΔE9 cells were pre-treated with 0, 2, or 5 mM MβCD for 30 min. Then, cells were washed and replenished with fresh culture media for 2 h. Following incubation, 1 ml of culture media was collected and centrifuged at 12,000 rpm for 5 min to spin down cell debris. Aβ42 levels were measured using a High Sensitivity Human Amyloid β42 ELISA Kit (Millipore, #EZHS42).

### Statistical analysis

Data are expressed as mean ± SEM. We conducted statistical analysis using one way ANOVA between the controls and the treated experimental groups; and considered *P* < 0.05 statistically significant.

## Acknowledgments

This work was supported by the Basic Science Research Program through the National Research Foundation of Korea funded by the Ministry of Education, Science and Technology (2016R1D1A1A099) to S.C.

## Authors’ contributions

YYC and SC designed the study. YYC and OHK performed cell cultures and all experiments. SC and YYC performed statistical analyses. SC and YYC wrote the manuscript with critical evaluation and comment by OHK. All authors read and approved the final manuscript.

## Additional Information

### Conflict of interests

The authors declare no competing financial interests.

## Expanded View figure legends

**Expanded View figure 1. APP localization in cholera toxin B positive-raft microdomains was altered by cellular cholesterol in CHO PS1 WT and PS1 ΔE9 cells**.

A. CHO PS1 WT cells were incubated with 150 μM MβCD-cholesterol and PS1 ΔE9 cells were treated with 5 mM MβCD. Then, cells were incubated with 6E10 antibody and 10 μg/ml of cholera toxin B (CTB) at 4°C to label surface APP and lipid raft microdomains, respectively. Representative confocal image demonstrated the co-localization of APP and CTB. Data are analyzed from five independent experiments. Scare bars correspond to 10 μm.

B, C, D. The ratio of overlap APP over CTB are indicated for (B) CHO PS1 WT cells and PS1 ΔE9 (n=5), (C) PS1 WT control and cholesterol-treated PS1 WT cells (n=5), and (D) PS1 ΔE9 and MβCD-treated PS1 ΔE9 cells (n=5). The co-localization of APP and CTB was determined with Image J.

Statistical analysis was carried out by one-way ANOVA: ***p<0.001.

**Expanded View figure 2. APP endocytosis rate in PS1 WT cells was increased by increasing cellular cholesterol levels**.

CHO PS1 WT cells were pre-treated with 150 μM MβCD-cholesterol to increase cellular cholesterol level. Cells were then labeled with APP antibody at 4°C to visualize APP endocytosis as described in Fig 2.

A. Representative confocal images show the localization of APP at indicated time points from four independent experiments. Scare bars correspond to 10 μm.

B. APP endocytosis was measured as the ratio of internalized APP over surface APP (n=4).

C. APP endocytosis was quantified using EZ-Link sulfo-NHS-SS-biotin as described in Fig 2. Total biotin-labeled APP (surface APP; sur.) was also detected. The upper panel shows a typical western blot result. The lower panel shows the rate of APP endocytosis by comparing internalized APP to surface APP (n=4). The band density was detected by a LAS-3000 system (Fuji Film, Japan) and analyzed with Multi Gauge software.

Statistical analysis was carried out by one-way ANOVA: ***p<0.001.

**Expanded View figure 3. Lipid raft fractionation**.

CHO PS1 WT and PS1 ΔE9 cells were collected with sodium carbonate buffer and fractionated with discontinuous sucrose density gradients as described in Methods. A total of 12 fractions were obtained from the top to the bottom, and equal volumes of each fraction were run on western blot to monitor the localization of total APP.

A, B. A typical western blot image indicated localization of APP, β-actin, and caveolin (lipid raft marker) (n=5) within 12 fractions (A) from PS1 WT and (B) PS1 ΔE9 cells, respectively.

C, D. (C) From PS1 WT and (D) PS1 ΔE9 cells, protein and cholesterol levels in each fraction were measured (n=5).

**Expanded View figure 4. Cholesterol affected accumulation of APP in early endosomes in CHO PS1 WT and CHO PS1 ΔE9 cells**.

A. CHO PS1WT cells were pre-treated with 150 μM MβCD-cholesterol. Cells were incubated with APP antibody (6E10) at 4°C and transferred to 37°C for indicated times to allow internalization. Following fixing and permeabilizing, cells were stained with EEA1 antibody to label early endosomes as described in Fig 6. Typical confocal image from four independent experiments shows co-localization of internalized APP and EEA1 at the indicated time points. Scare bars correspond to 10 μm.

B. CHO PS1 ΔE9 cells were pre-treated with 5 mM MβCD. Surface APP was labeled with 6E10 antibody at 4°C and internalized at 37°C for indicated times. Then, early endosomes were stained with EEA1 antibody after cells were fixed and permeabilized as described in Fig. 6. Representative confocal image from four independent experiments indicated APP localization within early endosomes. Scare bars correspond to 10 μm.

**Expanded View figure 5. Modulating cellular cholesterol levels regulated secreted Aβ42 levels in CHO PS1 WT and PS1 ΔE9 cells**.

A. Aβ42 levels were measured from the conditioned media in CHO PS1 WT and PS1 ΔE9 cells using Aβ42 specific ELISA kit (n=5).

B. CHO PS1 WT cells were incubated with 0, 75, or 150 μM MβCD-cholesterol. Then, cells were washed and replenished with conditioned media for 2 h. Aβ42 levels were analyzed from the conditioned media (n=4).

C. CHO PS1 ΔE9 cells were pre-treated with 0, 2, or 5 mM MβCD, and then cells were refreshed with new conditioned media for 2 h. Levels of Aβ42 were measured from conditioned media (n=5). Statistical analysis was performed by one-way ANOVA: **p<0.01, ***p<0.001.

## Appendix figure legend

**Appendix figure S1. Cellular cholesterol levels in CHO PS1 WT and PS1 ΔE9 cells**.

A. Total membrane cholesterol was measured in CHO PS1 WT and PS1 ΔE9 cells (n=7) using Amplex Red Cholesterol Assay Kit (Invitrogen, #A12216).

B. CHO PS1 WT cells were pre-treated with 0, 75, or 150 μM MβCD-cholesterol to increase cellular cholesterol levels (n=6), and (c) CHO PS1 ΔE9 cells were incubated with 0, 2, or 5 mM MβCD to decrease cellular cholesterol levels (n=6).

Statistical analysis was carried out by one-way ANOVA: *p<0.05, **p<0.01, ***p<0.001.

**Appendix figure S2. The endocytosis rate of transferrin was not affected by elevated cellular cholesterol levels**.

A. Cells were treated with Alexa488-conjugated transferrin in PBS at 37°C for indicated time periods to permit endocytosis. After removing remaining transferrin with acidic buffer, cells were fixed and visualized under a fluorescence microscope. Representative confocal image was from three independent experiments. Scare bars correspond to 10 μm.

B. Fluorescence intensities of internalized transferrin were analyzed using Image J software (n=3).

C. Cells were incubated with Alexa488-conjugated transferrin at 37°C for varying time periods to allow internalization. After washing, cells were harvested, and the same amount of protein was run on western blots (n=4).

D. Fluorescent bands were detected by a LAS-3000 system (Fuji Film, Japan) and were analyzed with the Multi Gauge software (n=4).

**Appendix figure S3. Surface APP localization in lipid raft microdomains was affected by the cholesterol levels in HeLa APPswe cells**.

Surface proteins were labeled with biotin at 4°C before fractionation. Lipid rafts (4 to 6, R) or non-lipid rafts (8 to 12, NR) were obtained as described in Fig 4. Then, the same amount of biotin-labeled proteins was captured with streptavidin beads, and captured biotin-labeled proteins were run on western blots to detect APP, β-actin, GAPDH, flotillin, and caveolin.

A. HeLa APPswe cells were pre-treated with 150 μM MβCD-cholesterol before labeling all surface proteins with EZ-Link NHS-biotin. The representative western blot image indicates the localization of surface APP in lipid raft and non-lipid raft fractions (n=5).

B. Relative band densities show the ratio of surface APP and internalized labeled APP from raft and non-raft fractions (n=5).

C. HeLa APPswe cells were pre-incubated with 1 mM MβCD before labeling all surface proteins and fractionation as described in Fig 4. Representative western blot result demonstrated biotin-labeled surface APP localization from lipid raft and non-lipid raft fractions (n=5).

D. The ratio of band intensity (n=5).

Statistical analysis was carried out by one-way ANOVA: *p<0.05.

**Appendix figure S4. Cellular cholesterol levels altered raft-dependent APP endocytosis in HeLa APPswe cells**.

T To verify raft-induced APP endocytosis, cells were biotinylated at 4°C to label surface proteins. Then, cells were transferred to 37°C for 10 min to allow internalization of biotin-labeled surface proteins, followed by fractionation as described in Fig 5. After fractionation, the equal amount of biotin-labeled protein from fractions 4-6 (lipid rafts, R) or 8-12 (non-lipid rafts, NR) was captured with streptavidin beads. Internalized biotin-labeled proteins were run on western blotting.

A. HeLa APPswe cells were pre-treated with 150 μM MβCD-cholesterol before fractionation. The corresponding western blot result shows the localization of biotin-labeled internalized APP (n=5).

B. The relative band density indicates the ratio of internalized APP from lipid raft and non-lipid raft fractions (n=5).

C. HeLa APPswe cells were incubated with 1 mM MβCD. Before fractionation, all surface proteins were labeled and internalized for 10 min as described in Fig 5. Representative western blot results demonstrate internalized APP levels from raft and non-raft fractions (n=4).

D. The band densitometry of internalized APP in lipid raft and non-lipid raft fractions was analyzed (n=4).

Statistical analysis was performed by one-way ANOVA: *p<0.05.

